# EphB6 regulates TFEB-lysosomal pathway and survival of disseminated quiescent breast cancer cells

**DOI:** 10.1101/2020.11.10.376186

**Authors:** Manuela Zangrossi, Probir Chakravarty, Patrizia Romani, Colin D.H. Ratcliffe, Steven Hooper, Martina Dori, Mattia Forcato, Silvio Bicciato, Sirio Dupont, Erik Sahai, Marco Montagner

## Abstract

Late relapse of disseminated cancer cells is a common feature of some types of tumors. Several intrinsic and extrinsic factors have been shown to affect reawakening of disseminated dormant cancer cells (DDCCs); however, the signals and processes sustaining survival of DDCCs in a foreign environment are still poorly understood. We have recently shown that crosstalk with lung epithelial cells promotes persistence of DDCCs from estrogen receptor positive (ER+) breast tumors. Here we show that TFEB-lysosomal axis is activated in DDCCs and that it is modulated by the pro-survival ephrin receptor EphB6. TFEB lysosomal direct targets are enriched in DDCCs *in vivo* and correlate with relapse in ER+ breast cancer patients. Direct contact of DDCCs with alveolar type I-like lung epithelial cells drives lysosomal accumulation and EphB6 induction. EphB6 contributes to TFEB transcriptional activity and lysosome formation in DDCCs *in vitro* and *in vivo*, and supports survival of DDCCs in coculture and *in vivo*. Furthermore, signaling from EphB6 promotes the proliferative response of surrounding lung parenchymal cells *in vivo*.

## Introduction

The time required to form overt metastases upon dissemination to a secondary organ varies considerably according to tissue of origin and subtype of the tumor (Risson et al. 2020; Klein 2020). Estrogen receptor positive breast cancers are amongst those cancer types whose latency period can reach 15 years (Zhang et al. 2013; Pantel and Hayes 2018), leading to the question of how DDCCs manage to survive in a foreign environment and reawaken after such a long time. Intertwined cell-intrinsic (Gawrzak et al. 2018; Pavlovic et al. 2015) and cell-extrinsic mechanisms sustain adaptation and exit from quiescence of DDCCs in the foreign environment (Aguirre-Ghiso and Sosa 2018). Inflammation (De Cock et al. 2016; Albrengues et al. 2018), stromal cells (Montagner et al. 2020; Malladi et al. 2016; Lu et al. 2011; Chen et al. 2011; Sosnoski et al. 2015; Ghajar et al. 2013; Wang et al. 2015; Ombrato and Montagner 2020), extracellular matrix proteins and architecture (Barkan et al. 2010; Ghajar et al. 2013; Montagner et al. 2020; Montagner and Dupont 2020; Montagner and Sahai 2020; Malanchi et al. 2011; Lowy and Oskarsson 2015; Albrengues et al. 2018), diffusible ligands (TGFβ1 (Ghajar et al. 2013), BMP (Gao et al. 2012), Wnt (Oskarsson et al. 2011), Notch (Capulli et al. 2019)) and hypoxia (Johnson et al. 2016; Fluegen et al. 2017) have been shown to regulate intracellular sensors that drive the choice between sustained quiescence and proliferation of DDCCs of breast origin (P-ERK/P-p38 ratio (Aguirre-Ghiso and Sosa 2018); PI3K/Akt/mTOR pathway (Chen et al. 2011; Wang et al. 2015; Balz et al. 2012); integrins (Montagner et al. 2020; Barkan et al. 2008; Touny et al. 2014; Albrengues et al. 2018; Barney et al. 2019; Carlson et al. 2019)). However, the mechanisms supporting survival of DDCCs, as opposed to those triggering exit from quiescence, are still unknown.

Because of the inherent asymptomatic nature of this process, isolation of DDCCs from healthy patients is often technically and ethically not possible, thus, we and others developed *in vitro* systems to study DDCCs-stroma crosstalk (Montagner and Sahai 2020; Montagner et al. 2020). A lung organotypic system, employing a defined combination of lung epithelial cells and fibroblasts, allowed us to recapitulate *in vitro* important features observed in DDCCs *in vivo*, such as Sfrp2-dependent development of filopodia-like structures, fibronectin fibrillogenesis and ultimately survival. Importantly, we showed that pro-survival and growth-restrictive signals emanating from alveolar type I cells (AT1) coexist and are modulated by surrounding stromal cell types and biochemical environment. In the view of a therapy that kills DDCCs before their reawakening, the imperative is targeting pro-survival signals, thus we concentrated on the crosstalk between indolent cancer cells and AT1 cells in a condition of low nutrient medium (mitogen low nutrient low, MLNL), where those signals dominate.

## Results and Discussion

To understand the crosstalk between DDCCs and lung epithelial cells more fully, we performed transcriptomic profiling and Gene Set Enrichment Analysis (GSEA) of AT1-like lung epithelial cells and D2.0R breast cancer cells (a cellular model of DDCCs) in monoculture and co-culture. We additionally compared these analyses with RNA sequencing of DDCCs from lungs (Montagner et al. 2020). Remarkably, a highly significant number of pathways and genes were similarly regulated in D2.0R *in vivo* (lung-disseminated) and in coculture with respect to monoculture (Fig. 1A and Supplemental Fig. S1, Pearson correlations equal to 0.811 and 0.559, respectively). Importantly, our transcriptomic analysis also captured stromal responses that were previously observed *in vivo*, such as proliferation of AT1 cells (Fig.1B), indicating that our coculture faithfully models processes observed *in vivo* both in DDCCs and stromal cells.

**Figure 1.**
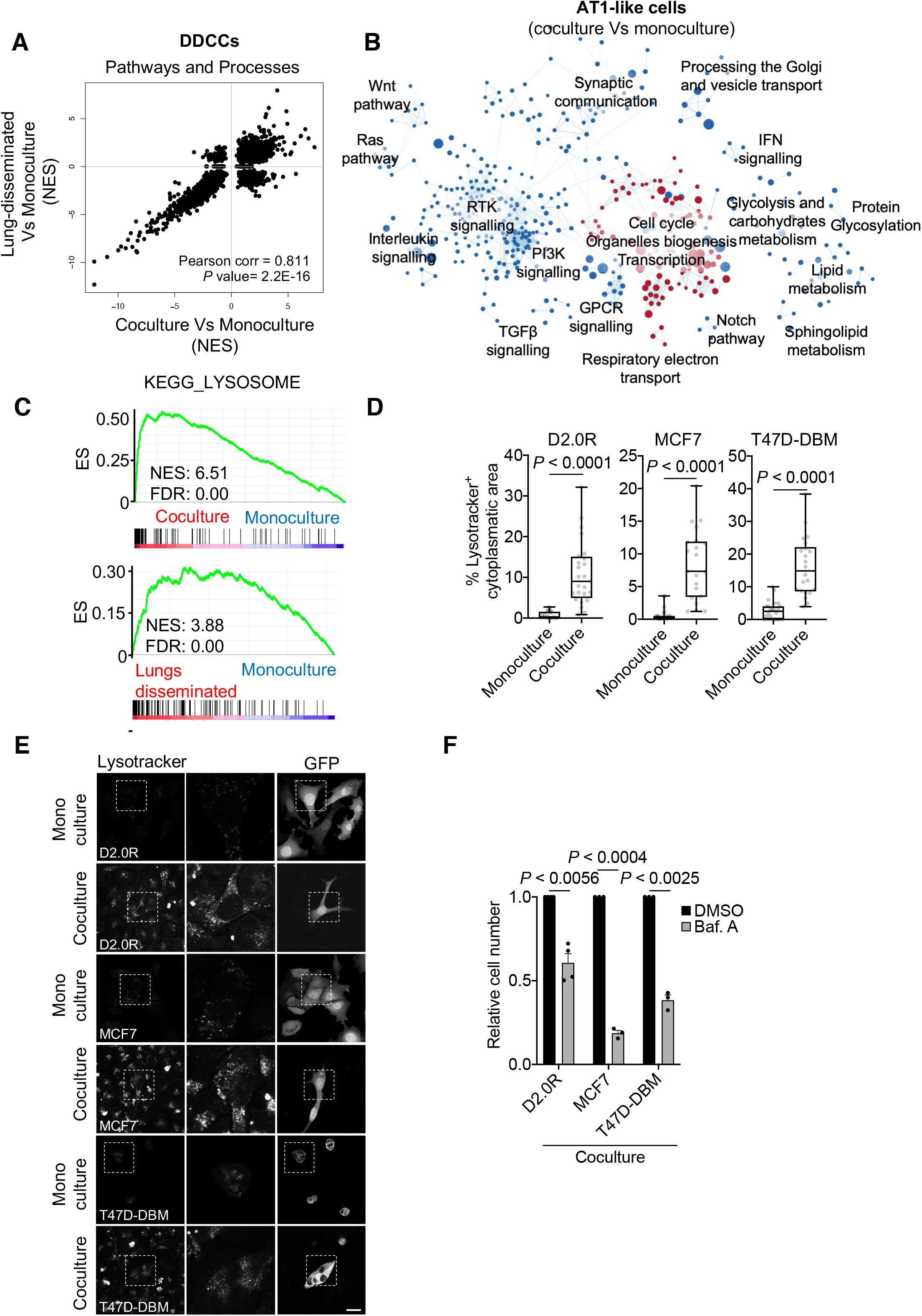
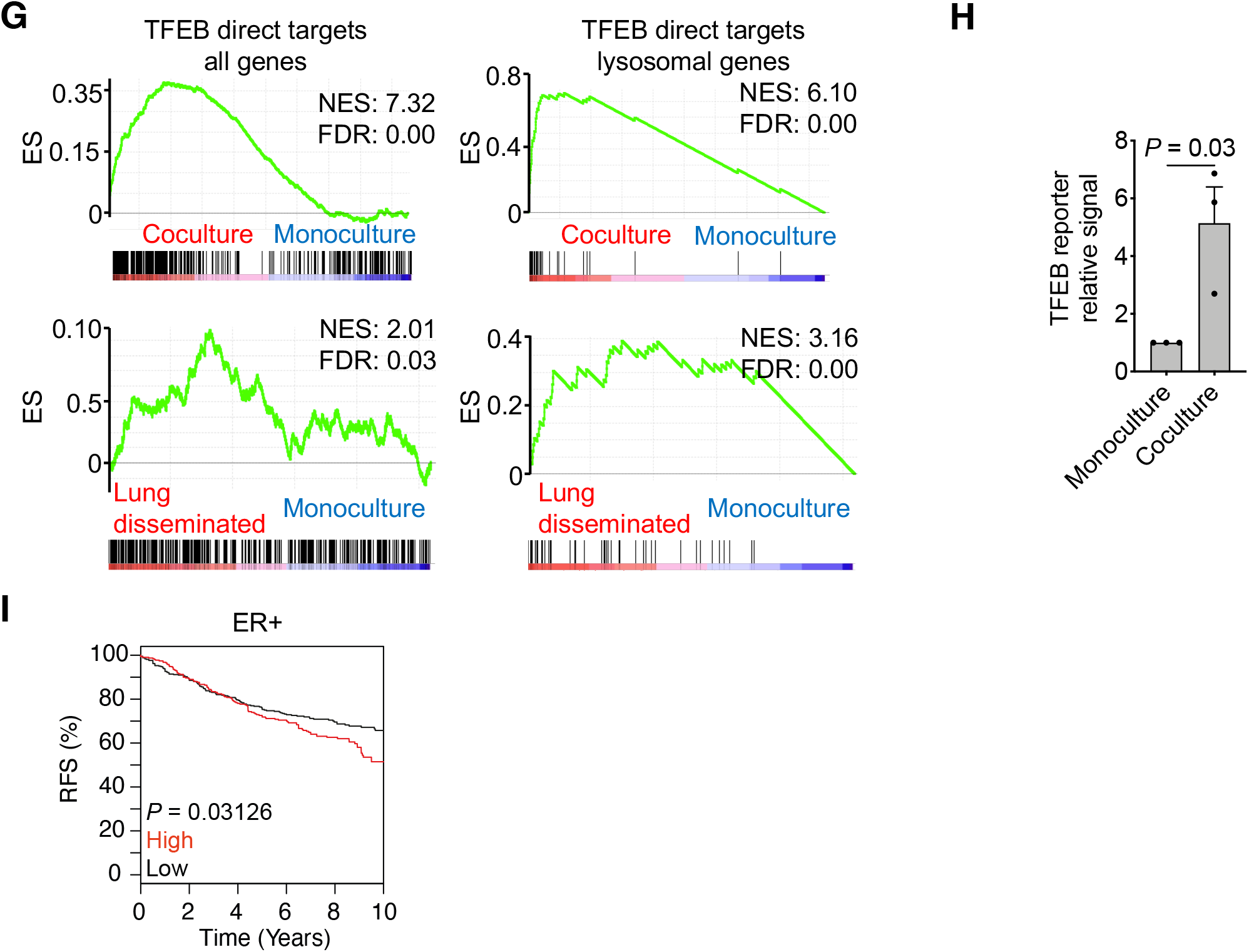
Disseminated dormant breast cancer cells activate TFEB-lysosomal axis in vivo and in lung-coculture. A. Scatterplot shows the correlation between the Normalized Enrichment Score of genesets included in Gene Set Enrichment Analysis (GSEA) of D2.0R-EGFP cells disseminated in vivo (lungs), in coculture or monoculture as indicated. B. Enrichment map for lung AT1-like cells upon coculture with indolent breast cancer cells. The map shows gene-set enrichment results of AT1-like cells cocultured with D2.0R-EGFP cells compared with AT1-like cells in monoculture. Node size, genes in pathway; node colour, enrichment score (red indicates enrichment in cocultured AT1-like cells, blue indicates enrichment monocultures of AT1-like cells); edge width, overlap size between connected nodes. C. Profile of the running ES score for KEGG_LYSOSOME geneset after GSEA of D2.0R-EGFP cells disseminated in vivo, in coculture or monoculture as indicated. D. Percentage of the cytoplasm with positive Lysotracker signal in the indicated indolent breast cancer cells lines upon monoculture or coculture with AT1-like cells. n=20-24 cells for D2.0R; n=21-29 cells for MCF7-EGFP; n=22-24 cells for T47D-DBM-EGFP. Mann-Whitney test. Whisker plot: midline, median; box, 25–75th percentile; whisker, minimum to maximum. E. Representative images of Lysotracker staining quantified in D. Dashed box: area magnified in the middle images. Scale bar: 20 μm. F. Relative cell number of the indicated indolent breast cancer cells upon coculture with AT1-like cells, after treatment with 4 μM Bafilomycin A for 4 days. n=4 for D2.0R, n=3 for MCF7 and T47D-DBM cells (independent experiments). Paired two tailed t-test, mean with SEM. G. Profile of the running ES score for genesets including TFEB direct targets (all targets or subselection of lysosomal genes, as in (Palmieri et al. 2011)) after GSEA of D2.0R-EGFP cells disseminated in vivo, in coculture or monoculture as indicated. H. Relative induction of transfected TFEB-luciferase reporter in D2.0R-EGFP cells cocultured with AT1-like cells compared to monoculture. n=3 independent experiments, ratio paired two-tailed t-test, mean with SEM. I. Kaplan-Meier curve showing Relapse-Free Survival of ER+ breast cancer patients derived from the database at http://co.bmc.lu.se/gobo/gsa.pl, stratified according to the TFEB signature.

Gene signatures related to lysosomal biogenesis and vesicle transport stood out as a process significantly upregulated after dissemination and upon coculture with AT1-like cells (Fig. 1C). In line with this finding, lysosome accumulation was observed in different DDCCs in coculture compared to monoculture, as visualized by staining with Lysotracker, a specific dye for acidic organelles (Fig. 1D and E). Having established an *in vitro* system that mimics lung-specific *in vivo* processes upon dissemination of DDCCs, we tested if functional lysosomes are required for survival of DDCCs in this context. Importantly, blocking lysosome acidification with bafilomycin A1 led to reduced surviving DDCCs in the coculture without affecting stromal cells (Fig. 1F and Supplemental Fig. S2).

As TFEB, a member of MiT-TFE family of transcription factors, is a master regulator of lysosome biogenesis and associated metabolic processes (Napolitano and Ballabio 2016; Lawrence and Zoncu 2019; Perera and Zoncu 2016), we queried our transcriptomic analysis with a gene list of 471 TFEB direct targets and with a selection of TFEB direct targets with known lysosomal function (Sardiello et al. 2009; Palmieri et al. 2011). Both signatures were highly enriched in DDCCs *in vivo* (lung-disseminated) and cocultured dormant cells, suggesting that activated TFEB might be responsible for the observed lysosomal accumulation (Fig. 1G). TFEB transcriptional activation upon coculture was confirmed by a TFEB luciferase reporter in DDCCs (Fig. 1H). Interestingly, higher expression of TFEB direct targets in ER+ breast cancer patients was linked with increased number of relapses at delayed time points (Fig. 1I). These data suggest the activation of TFEB-dependent lysosomal biogenesis in DDCCs *in vivo* and in a lung coculture system and support its involvement in survival of disseminated indolent breast cancer cells.

We then sought to identify mediators of DDCCs-lung stroma crosstalk. We first asked whether direct contact among the different cell types was required for activation of the dormant phenotype. Quantitative PCR with reverse transcription (RT-qPCR) for representative dormancy-associated genes (Montagner et al. 2020) revealed that conditioned medium from AT1-like cells was not sufficient to trigger transcription of genes that are instead induced by cell-cell contacts following direct coculture (Fig. 2A). We thus focused on cell surface signaling molecules that might be involved in communication between lung epithelial cells and indolent breast cancer cells. An *in vivo* loss-of-function screen identified genes required for the survival of breast DDCCs, such as *Sfrp2*, i.e. genes whose depletion caused death of DDCCs upon dissemination to the lungs (Montagner et al. 2020). Among those genes we noticed *EphB6*, a member of the Eph family of receptor tyrosine kinase (Fig. 2B). Interestingly, EphB receptors’ ligands, ephrin-Bs, are transmembrane proteins and are thus a good candidate to explain contact-mediated processes (Kania and Klein 2016; Liang et al. 2019; Nikas et al. 2018). Importantly, we repeated the results from the screening with multiple short hairpin RNA targeting *EphB6*, validating its relevance in the context of persistence of DDCCs (Fig. 2C).

**Figure 2.**
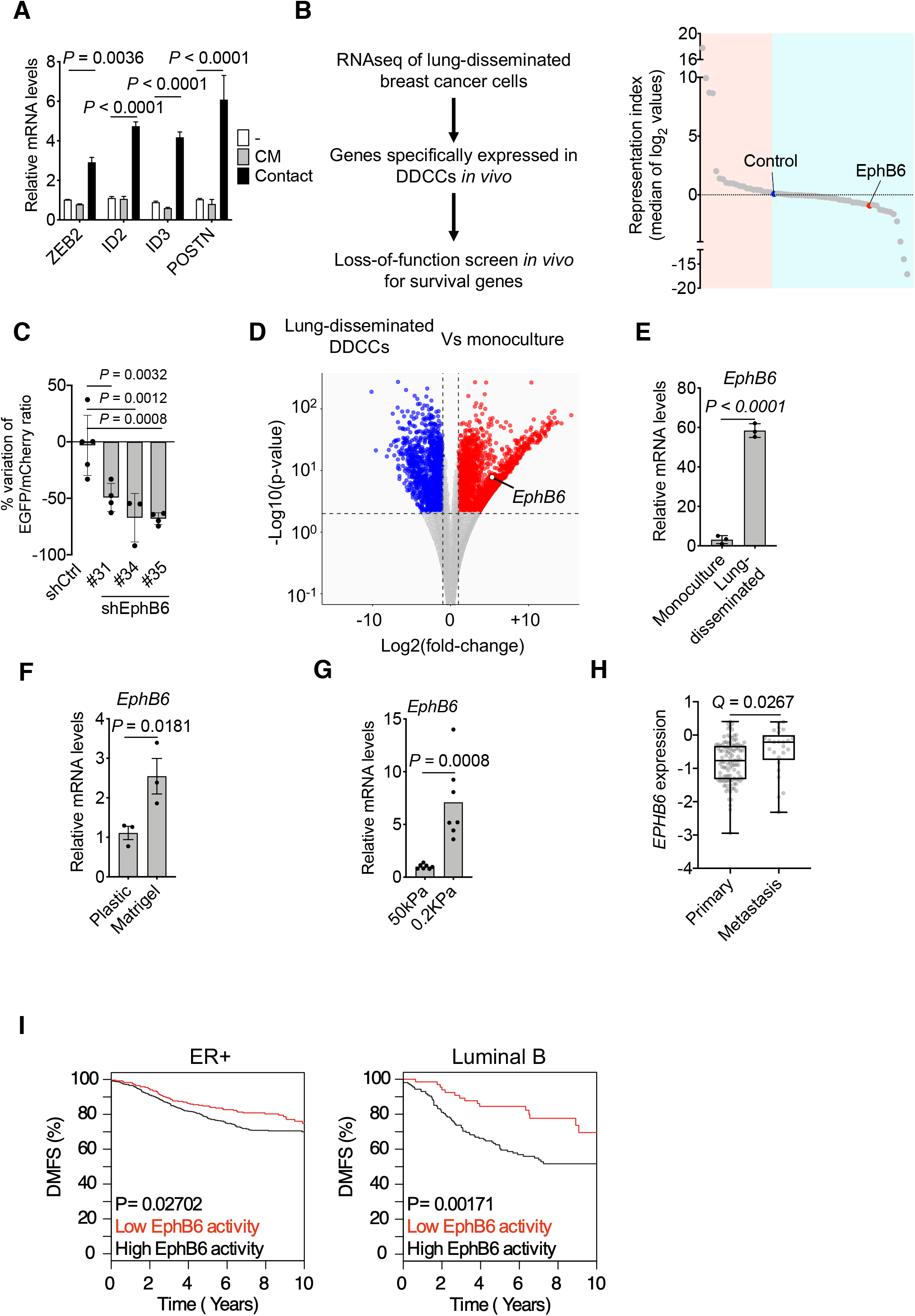
EphB6 supports survival of disseminated dormant breast cancer cells. A. RT-qPCR analysis of representative genes of the dormancy response. D2.0R-EGFP cells have been cultured alone, cocoltured with AT1-like cells (Contact) or with conditioned medium from AT1-like cells (CM). n=3. Two-way ANOVA test, multiple comparisons. Mean with SEM. B. Left, workflow of loss-of-function screen in vivo for survival genes in DDCCs (Montagner et al. 2020). Right, Representation scores for each gene included in the screen, calculated from the fold change of representation of each shRNA relative to pre-injection abundance. On the light blue side, there are genes whose downregulation leads to reduced representation of the clones, such as EphB6. C. D2.0R-EGFP cells stably expressing the indicating shRNA were injected intravenously together with an equal amount of D2.0R-mCherry-shCtrl cells as internal control. After 3 weeks the amount of surviving D2.0R cells was measured and the ratio EGFP/mCherry calculated. n=5 mice for shCtrl cells, n=4 mice for each shEphB6 sequence. One-way ANOVA test. Mean with SD. D. Transcriptome analysis (RNA sequencing) of D2.0R-EGFP cells upon dissemination in the lung Vs D2.0R-EGFP in monoculture. DOWN-regulated (Log2 fold-change < −1 and adjusted p-value < 0.01) and UP-regulated (Log2 fold-change > 1 and adjusted p-value < 0.01) genes are indicated in blue and red respectively. E. qPCR of EphB6 gene in D2.0R-EGFP cells upon dissemination in the lung (n=3 mice) or after monoculture (n=3 samples). Unpaired t-test. Mean with SD. F. Relative expression of EphB6 gene in D2.0R-EGFP cells cultivated on coated plastic or on Matrigel. n=3 independent experiments, ratio paired two-tailed t-test, mean with SEM. G. Relative expression of EphB6 gene in D2.0R-EGFP cells cultivated on synthetic hydrogels with indicated stiffness. n=7 samples merged from n=3 independent experiments, unpaired two-tailed t-test. H. EphB6 expression in ER+ primary breast cancers and metastases from publicly available databases (Harrell et al. 2012). q-value after unpaired Significance Analysis of Microarray. I. Kaplan-Meier curves showing Distant Metastasis-Free Survival of indicated breast cancer patients with indicated cancer subtypes derived from the database at http://co.bmc.lu.se/gobo/gsa.pl, stratified according to genes repressed by EphB6. Black line indicates patients with lower expression of those genes, i.e. with higher EphB6 activity, that is correlated to increased likelihood of distant relapses.

An additional feature pointing to a role for EphB6 in the communication between lung epithelial cells and DDCCs was the observation that *EphB6* mRNA was upregulated in lung-disseminated DDCCs compared to culture on plastic (Fig. 2D and 2E) in indolent breast cancer cells. As lung parenchyma is characterized by an ECM with low stiffness (Young’s modulus of approximately 0,5-2 kPa according to (Booth et al. 2012; Liu et al. 2010)), we hypothesized that a soft microenvironment could contribute to *EphB6* induction in DDCCs. We tested this hypothesis by assessing *EphB6* expression in indolent breast cancer cells cultivated on substrates with different stiffness. First, *EphB6* was induced when cells are cultured on top of a soft naturally-derived 3D ECM scaffold (Matrigel), compared to ECM-coated stiff plastic substrate (Fig. 2F). Second, we cultivated D2.0R cells on synthetic ECM-coated acrylamide hydrogels of controlled stiffness and confirmed *EphB6* induction at low Young’s modulus values (Fig. 2G). Notably, *EphB6* was also found upregulated in breast cancer cells from metastases compared to estrogen receptor positive primary breast cancers (Fig. 2H). Next, we investigated the link between EphB6-dependent gene program and human breast cancer. A signature of genes upregulated in *EphB6*-depleted cells (i.e. genes reflecting a low EphB6 activity) were associated with improved distant metastasis-free survival (DMSF) of ER+ subtypes of breast cancers (Fig. 2I). These results support a model whereby EphB6 is induced *in vivo* in soft microenvironments and has a role in the survival of indolent disseminated breast cancer cells.

Eph-ephrin stimulation is bidirectional and signals are propagated in Eph-expressing cells as well as in ephrin-expressing cells (forward and reverse signaling, respectively (Kania and Klein 2016; Liang et al. 2019; Nikas et al. 2018)). We then asked whether EphB6 expression in breast cancer cells could influence neighboring AT1 cells. RNA sequencing of cocultured AT1-like cells revealed that depletion of EphB6 in D2.0R cells led to downregulation of several cell cycle-related pathways and upregulation of metabolic and other signaling pathways (Fig. 3A). In agreement with this, less proliferating lung stromal cells were observed in the proximity of EphB6-deficient DDCCs *in vivo* (Fig. 3B and C). We then turned our attention to the role of EphB6 in DDCCs and compared at genome-wide level, transcriptomes of EphB6-depleted DDCCs cells in coculture and *in vivo*. Pearson correlation coefficients highlighted that a large number of pathways and processes are affected by EphB6 depletion (with two independent short interfering RNAs) both *in vivo* and in coculture compared to control cells (Fig. 3D). Strikingly, lysosomal and other vesicle biogenesis signatures were amongst the processes most strongly affected by EphB6 knockdown in coculture and *in vivo* (Fig. 3E). In line with our findings on the role of TFEB and lysosomal biogenesis in DDCCs shown above, we found that TFEB transcriptional activity was significantly reduced in cells with short interfering RNAs against EphB6, suggesting a connection between EphB6 and TFEB (Fig. 3F).

**Figure 3.**
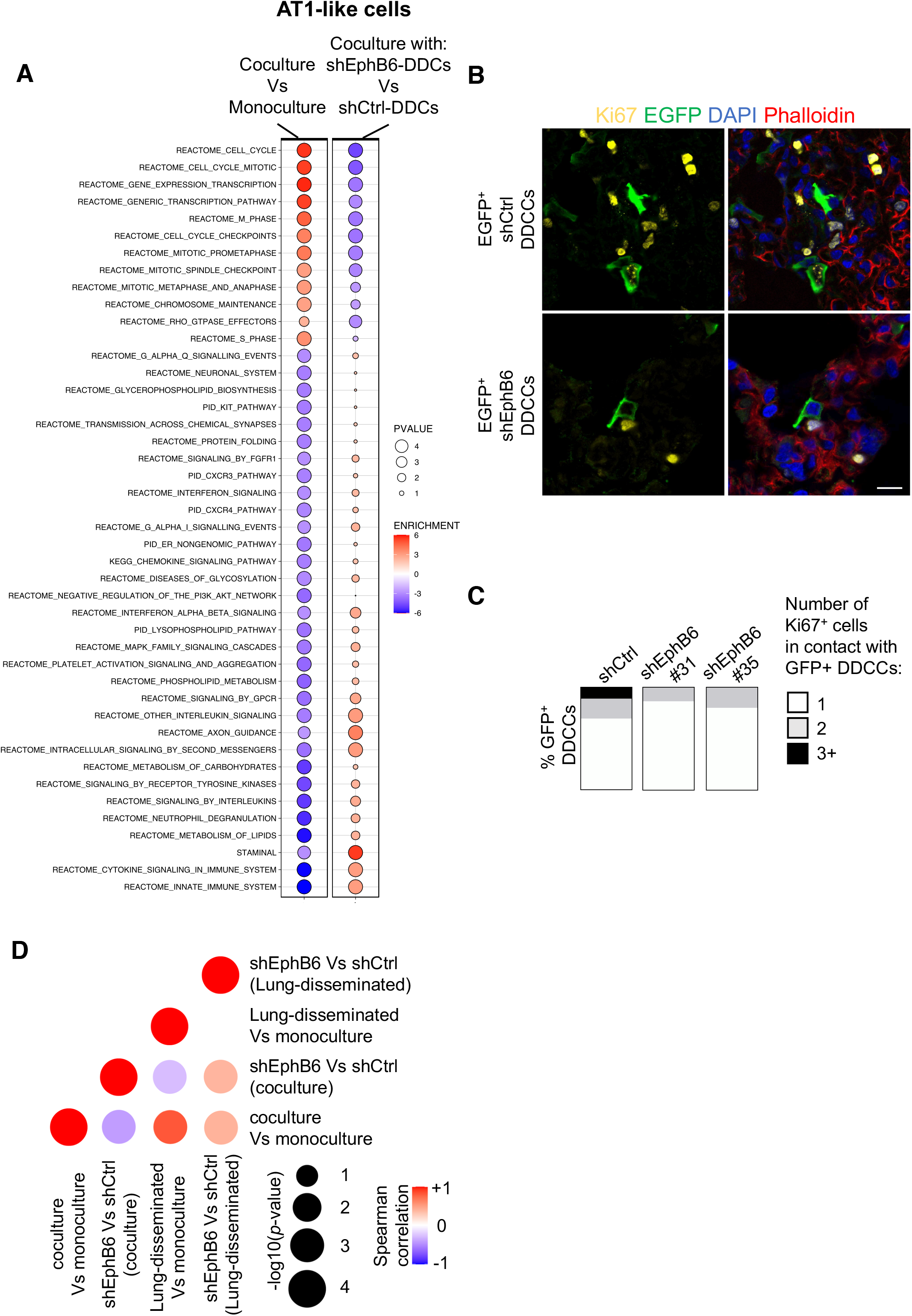
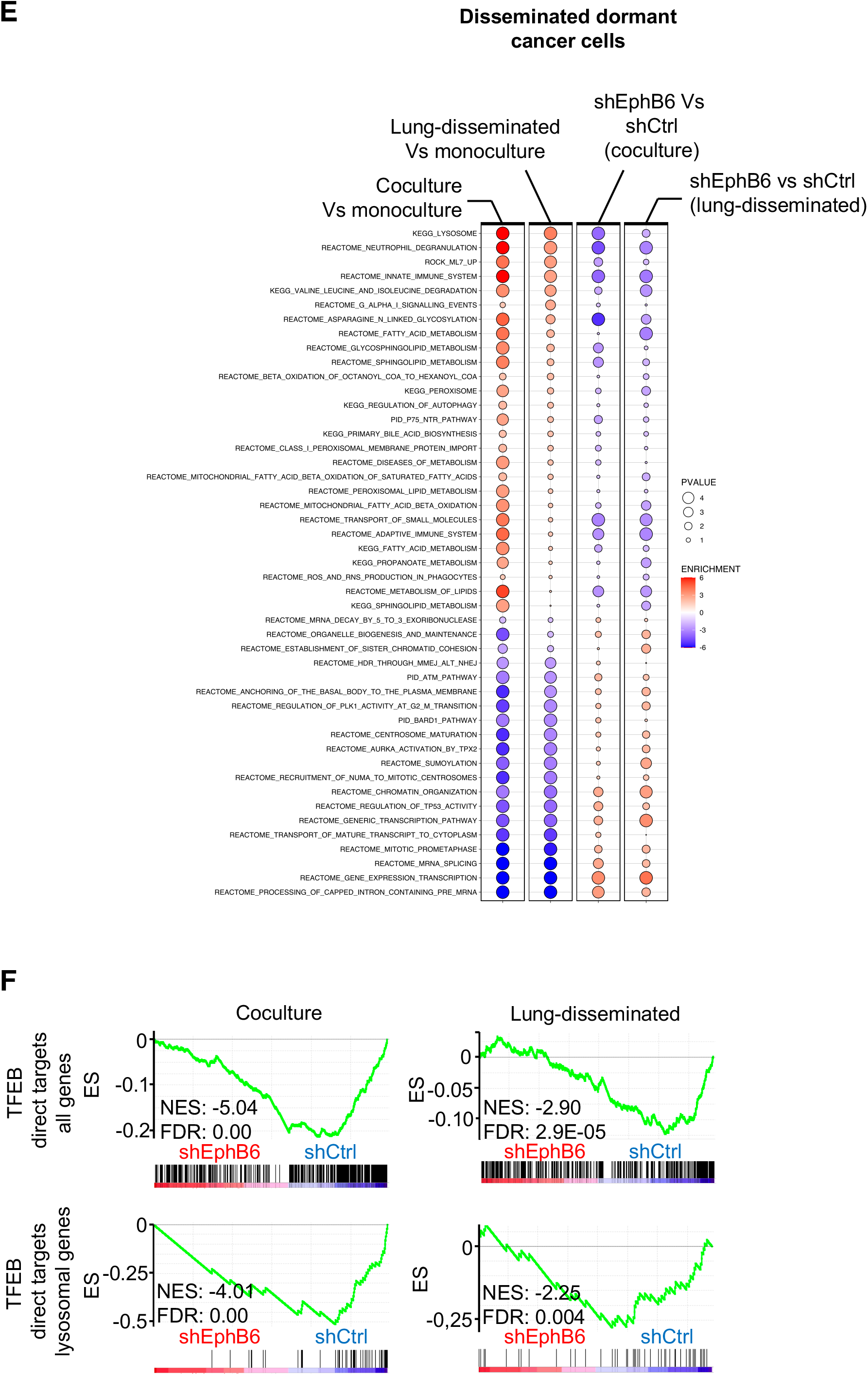
EphB6 is required for the transcriptional responses of disseminated dormant breast cancer cells and lung stromal cells. A. Balloon plot summarizing GSEA results of AT1-like cells in monoculture and cocultured with D2.0R-EGFP-shCtrl cells or shEphB6 cells (results from two independent shEphB6 sequences, #31 and #35). Balloon size represents the statistical significance (−log10 FDR), while colour indicates the fold-enrichment for each term (NES). B. Representative images of Ki67^+^ lung stromal cells surrounding GFP^+^ disseminated indolent breast cancer cells with control shRNA or shRNAs targeting EphB6. Scale bar, 10 μm. C. Quantification of images in B. Percentage of D2.0R-EGFP cells in contact with the indicated Ki67^+^ stromal cells. n = 165, 119 and 147 cells for shCtrl, shEphB6#31 and shEphB6#35, respectively, merged from 3 mice/sample. Data showed as parts of a whole. D. Dot plot shows the correlation of NES values generated from GSEA between four indicated comparisons, where the color represents the Spearman correlation and size presents the −log10(p-value) of the correlation. E. Balloon plot summarizing GSEA results of the indicated comparisons for each indicated genesets. The plot was manually curated to help visualization and shows genesets with higher coherent enrichment in the different conditions. Balloon size represents the statistical significance (−log10 FDR), while colour indicates the fold-enrichment for each term. F. Profile of the running ES score for genesets including TFEB direct targets (all targets or subselection of lysosomal genes) after GSEA of D2.0R-EGFP cells with shCtrl or shEphB6 either disseminated in vivo or in coculture with AT1-like cells.

To better understand the link between EphB6 and lysosomal biogenesis, we exploited our coculture system. We showed that EphB6 knockdown decreases TFEB transcriptional activity (Fig. 4A) and consequently lysosomal accumulation in coculture and *in vivo* (Fig. 4B-D). Importantly, EphB6 is required for the survival of indolent breast cancer cells cultivated together with lung epithelial cells (Fig. 4E), as previously observed *in vivo* (Fig. 2C). Collectively this data suggest that downregulation of EphB6 affects TFEB transcriptional activity, lysosome accumulation and survival of DDCCs.

**Figure 4.**
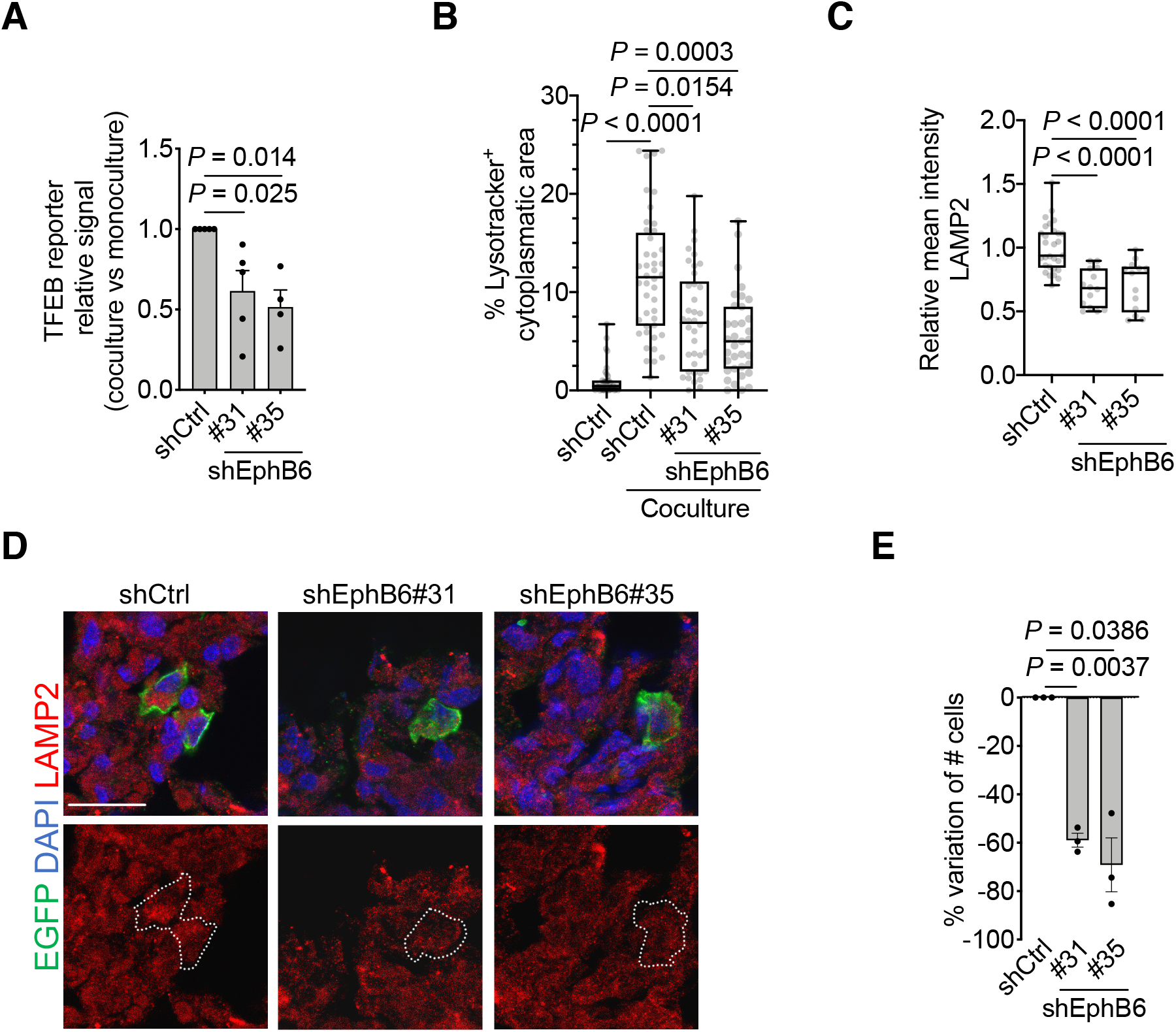
EphB6 regulates TFEB-lysosomal axis. A. Relative induction (coculture with AT1-like cells Vs monoculture) of transfected TFEB-luciferase reporter in D2.0R-EGFP cells stably expressing the indicated shRNAs. n=5 (shCtrl and shEphB6#31) or 4 (shEphB6#35) independent experiments. Dunn’s multiple comparisons test. Mean with SEM. B. Percentage of the cytoplasm with positive Lysotracker signal in D2.0R-EGFP cells with indicated shRNAs upon coculture with AT1-like cells, or in monoculture. n = 46 cells for shCtrl on monoculture; n = 44 cells for shCtrl on coculture; n = 38 cells for shEphB6#31; n = 35 cells for shEphB6#35. Dunn’s multiple comparisons test. Whisker plot: midline, median; box, 25–75th percentile; whisker, minimum to maximum. C. Relative mean intensity of lung disseminated shCtrl-, or shEphB6-, D2.0R-EGFP cells. n = 29 cells for shCtrl, n = 13 or 14 cells for shEphB6 across 2 mice/sample. One-way ANOVA. Whisker plot: midline, median; box, 25–75th percentile; whisker, minimum to maximum. D. Representative images of lysosomal membrane protein LAMP2 in control D2.0R-EGFP cells or cells with EphB6 knockdown after dissemination to lung parenchyma quantified in C. Scale bar, 20 μm. E. Variation of cell number of D2.0R-EGFP upon shEphB6 knock-down in coculture with AT1-like cells. n = 3 independent experiments. One-way ANOVA. Mean with SEM.

As the leading cause of cancer-related death, the metastatic process has been the object of intense research in the last decades. However, effective prevention or metastases-specific therapies are still an elusive goal. Metastatic dormancy offers a therapeutic window so far unexploited, and yet processes associated with persistence of DDCCs are still largely unknown (Risson et al. 2020; Klein 2020; Massagué and Obenauf 2016). Our work suggests that EphB6 plays a critical role in the crosstalk of indolent breast cancer cells with alveolar type I and supports survival of DDCCs *in vivo* and *in vitro*. Interestingly, it has been reported that restoration of EphB6 expression in EphB6-null triple negative breast cancer cells (TNBC), induced cell spreading, actin-dependent protrusions and scattered proliferation of tumor initiating cells (Truitt et al. 2010; Toosi et al. 2018), in agreement with some features observed in ER+ DDCCs. *EphB6* mRNA level is higher in soft natural and synthetic scaffolds as well as in metastatic lesions (Fig. 2F-H). Moreover, *EphB6* mRNA expression depends also on ECM composition, in addition to stiffness, as evidenced in publicly available RNA sequencing analysis of cells plated on collagenI-enriched Matrigel (Vera-Ramirez et al. 2018). This observation may suggest that disseminated clones with higher EphB6 expression, having increased fitness, might participate in metastatic outgrowth. In line with bidirectional Ephs-ephrins signaling, *EphB6* knock-down in DDCCs affected proliferation of neighboring AT1 cells *in vitro* and *in vivo* (Fig. 3A-C). EphB6 reduction is accompanied by reduced TFEB-dependent genes transcription in indolent breast cancer cells from lungs and lung-organotypic system (Fig. 3F) and decreased cytoplasmic area with Lysotracker-positive organelles (Fig. 4B).

Lysosomes are the cellular hub that integrates degradation/recycling of cellular components with stress responses allowing dynamic metabolic adaptation, an essential asset for cells disseminated in a foreign microenvironment. Trafficking routes that funnel into lysosomal degradation include clathrin-dependent and independent endocytosis, phagocytosis and macropinocytosis, macroautophagy, chaperone-mediated autophagy, integrin-mediated scavenging (Muranen et al. 2017) and entosis (Overholtzer et al. 2007). Interestingly, lysosome regulation has been found key in the modulation of quiescence, proliferation and differentiation of hematopoietic stem cells (Liang et al. 2020). Although the specific role of lysosomes has not been addressed before in metastatic dormancy, conflicting data on autophagy have been reported, with opposite phenotypes upon knock-down of different autophagy mediators (Vera-Ramirez et al. 2018; La Belle Flynn et al. 2019). Our data support the view of lysosomal flux as an essential process for the survival of DDCCs in the lung, however future investigations are required to understand whether this is due to specific cargoes converging on a greater number of lysosomes or a more general role for enhanced lysosomal-mediated turnover. Another important aspect that remains to be elucidated is whether this mechanism is shared among other tissues or other lung stromal cells. For example, *in vivo* labelling experiments revealed de-differentiation and proliferation of the lung-epithelial compartment (of AT2 origin) in the metastatic niche of aggressive lung-disseminated breast cancer cells (Ombrato et al. 2019). Whether EphB6 is involved in this specific crosstalk is an open question for future experiments. Data from our work describe a novel regulator of breast DDCCs survival, EphB6, that modulates adaptation to lung microenvironment through TFEB-lysosomal axis, providing potential novel liabilities of disseminated dormant breast cancer cells.

## Materials and Methods

### Cell lines

Alveolar type 1-like cells (TT1 cells) were a gift from J. Downward (The Francis Crick Institute, London) and were originally provided by T. Tetley (Imperial College, London). T47D-DBM cells were a gift from R. Gomis (Institute for Research in Biomedicine, Barcelona). D2.0R and MCF7-GFP cells were a gift from D. Barkan (University of Haifa). All cells were kept in DMEM with 10% FBS (Thermo Fisher Scientific, 41965-039) and routinely screened for mycoplasma at the Cell Services facility at The Francis Crick Institute or with Universal Mycoplasma Detection kit (ATCC, 30–1012 K).

### Stable protein expression and gene knock-down

Generation of D2.0R-EGFP and D2.0R-mCherry has been described in (Montagner et al. 2020). shRNA expressing cells were generated with lentiviral transduction. pLKO.1-based plasmids (MISSION, Sigma-Aldrich) were transfected into 293T cells together with packaging plasmids (pMD2, psPAX2). After 2 days, supernatants were collected, filtered through a 0.45 μm filter and added to indicated cells for 2 days before selection with puromycin. List of shRNA sequences is provided in Supplementary Table S1.

### Tissue dissociation

Lungs were dissociated to single-cell suspension as in (Montagner et al. 2020). Briefly, chopped lungs were digested with digestion solution (PBS buffer with 75 μg/ml TM Liberase (Roche, 05401151001), 75 μg/ml TH Liberase (Roche, 05401127001), 12.5 μg/ml DNAse (Sigma-Aldrich, DN25)) for 1 h at 37 °C on a rocker and homogenised by pipetting. After filtration to remove undigested clumps, red blood cells were lysed with Red Blood Cells Lysis Solution (Miltenyi Biotec, 130-094-183) following the manufacturer’s protocol. After washing, cells were resuspended in FACS buffer (PBS, 2 mM EDTA and 3% BSA) and labelled with CD45–APC antibody for 30 min (eBiosciences, 30-F11, 1:400) to avoid contamination from leukocytes during sorting.

### *In vivo* assays and quantifications

The study is compliant with all relevant ethical regulations regarding animal research. All protocols were in accordance with UK Home Office regulations under project license PPL80/2368 and subsequently PPL70/8380, which passed ethical review by the LRI Animal Welfare Ethical Review Board in 2014.

For tail vein injections, cells were resuspended in PBS and 150 μl/mouse injected using a 25G needle. At end point, mice were culled by a schedule 1 method. For quantification of disseminated indolent cells after EphB6 knockdown, 5 × 10^5^ D2.0R-mCherry shControl cells (Sigma-Aldrich, SHC016) were injected into the tail vein of 6- to 8-week-old female nude athymic BALB/c mice together with 5 × 10^5^ D2.0R-EGFP shControl cells or 5 × 10^5^ D2.0R-EGFP shEphB6. Lungs were collected and processed as in (Montagner et al. 2020). Number of CD45^−^/EGFP^+^ and CD45^−^/mCherry^+^ cells were quantified by FACS and the ratio EGFP/mCherry calculated to evaluate the survival of shRNA-bearing cells (EGFP) relative to an internal control (mCherry).

#### Sample preparation for RNA sequencing and qPCR analysis

D2.0R-EGFP-shControl (1×10^6^ cells/sample) or a mix of D2.0R-EGFP-shEphB6 (#31, #34, #35) were injected in the tail vein of nude athymic BALB/c mice. After two weeks, lungs were harvested and digested into a single-cell suspension as described above. CD45^−^/EGFP^+^ cells were sorted (Flow Cytometry Facility at Cancer Research UK-LRI and The Francis Crick Institute) directly into lysis buffer and total RNA was extracted with RNeasy Plus Micro Kit (Qiagen) following the manufacturer’s instructions.

### Lung organotypic system

#### Samples preparation for survival analysis and imaging

At day 1, 1,36×10^5^ TT1 cells/well were plated onto Lumox 24-multiwell plate (Sarstedt, 94.699.00.14) in MLNL medium (low-glucose DMEM/1% FCS, Thermo Fisher Scientific 21885025). At day 2, breast cancer cells (100 cells/well for survival assays, 500 cells/well for imaging) were plated onto TT1 cell layer in MLNL. For quantification, GFP^+^ cells were manually counted under an inverted fluorescent microscope after replacing the medium with HBSS.

#### Samples preparation for RNA sequencing

D2.0R-EGFP-shControl, D2.0R-EGFP-shEphB6#31 or D2.0R-EGFP-shEphB6#35 (1,8×10^5^ cells/sample) were plated as above and cocultured with AT1-like cells (1,36×10^6^ cells/sample) in MLNL medium (in 60 mm dish) for three days before separation by Fluorescence-Activated Cell Sorter (FACS) based on EGFP signal. Total RNA from three biological replicates/sample was extracted using RNeasy Plus Micro Kit (Qiagen). For bioinformatic analysis, replicate samples for D2.0R-EGFP-shEphB#31 and #35 were grouped together and compared against control sample to identify genes that were differentially expressed relative to control that were common between both shRNAs.

#### Samples preparation for qPCR analysis

Samples were prepared as for RNA sequencing, except that total RNA was extracted from the whole coculture, retrotranscribed, and mouse genes were amplified by using mouse-specific qPCR primers.

### Reverse Transcriptase Real Time PCR (RT-qPCR)

Total RNA was retrotranscribed with dT-primed M-MLV Reverse Transcriptase (Thermo Fisher Scientific, 28025013). qPCR analysis was carried out in a QuantStudio 6 Flex Real-Time PCR System (Thermo Fisher Scientific) with Fast SYBR Green Master Mix (Applied Biosystems 4385612). Gene expression values of EphB6 *in vivo* and cultivated on scaffolds were normalized to GAPDH. Gene expression values from cocultured Vs monocultured D2.0R cells were normalized to GFP expression levels (not expressed AT1-like cells). For RT–qPCR analysis of EphB6 gene in disseminated breast cancer cells *in vivo*, cells were isolated from lungs and total RNA was amplified with Arcturus RiboAmp HS PLUS kit to obtain enough cDNA for RT–qPCR analysis. List of primers used in qPCR is provided in Supplementary Table S2.

### Cell culture on natural and synthetic scaffolds

For experiments with natural scaffolds, wells were coated with 100% Matrigel (BD Bioscience) for soft substrate. For stiff substrate, 2% Matrigel was used to coat plastic for 1 hour and then removed.

For experiments with synthetic scaffolds, cells were plated on commercial soft (0.2 KPa, SOFTWELL SW12-COL-0.2 PK) or stiff hydrogels (50 KPa, SW12-COL-50 PK). Cells were harvested for total RNA isolation after 24 hours.

### Bioinformatics

#### RNA sequencing

Before analysis, RNA samples were assessed for quantity and integrity using the NanoDrop 8000 spectrophotometer v.2.0 (Thermo Fisher Scientific) and Agilent 2100 Bioanalyser (Agilent Technologies), respectively. Biological replicate libraries were prepared using the polyA KAPA mRNA HyperPrep Kit and sequenced on Illumina HiSeq 4000 platform, generating ~24 million 100bp single-end reads *per* sample. Read quality trimming and adaptor removal was carried out using Trimmomatic (version 0.36). The RSEM package (version 1.3.30)(Li and Dewey 2011) in conjunction with the STAR alignment algorithm (version 2.5.2a)(Dobin et al. 2013) was used for the mapping and subsequent gene-level counting of the sequenced reads with respect to Ensembl mouse GRCm.38.89 version transcriptome. Normalization of raw count data and differential expression analysis was performed with the DESeq2 package (version 1.18.1)(Love et al. 2014) within the R programming environment (version 3.4.3)([CSL STYLE ERROR: reference with no printed form.]). Differentially expressed genes were defined as those showing statistically significant differences (FDR <0.05). Differential gene lists ranked by the Wald statistic were used to look for pathway and selected genesets using the Broad’s GSEA software (version 2.1.0) with genesets from MSigDB (version 6)(Subramanian et al. 2005) and additional published and custom datasets (Supplementary Table S3). Spearman’s rank correlation was used to compare the normalized enrichment scores between comparisons from different experiments to determine which pathways were similarly enriched. Scatterplots (generated using the R base graphics package) shows the correlation between the Wald’s statistic (gene level differences from DESEQ2) or the normalized enrichment score (pathway level differences from GSEA) when comparing D2.0R lung-disseminated_Vs_monoculture and coculture_vs_monoculture comparisons. Dot plot (generated using R’s ggplot2 package) shows the correlation of NES values generated from GSEA between four indicated comparisons, where the color represents the Spearman correlation and size presents the −log10(*p*-value) of the correlation using the cor.test function. Volcano plot was produced using log2FC and adjusted p-value obtained by differential expression analysis exploiting the “ggscatter" function from ggpubr R package (v. 0.2). Balloon plots were made using “ggballoon” function from ggpubr R package (v. 0.2). For enrichment map, GSEA results from D2.0R versus other groups were visualized using Cytoscape (v.3.6.0) and the enrichment map plug-in (Merico et al. 2010). The map has been manually annotated to reduce complexity and redundancy.

#### Analysis of public datasets of primary and metastatic breast cancer samples

To gain insights into the expression of EphB6 in primary breast cancer and metastases, we analyzed publicly available data from microarray (GSE26338,(Harrell et al. 2012)). We downloaded from Gene Expression Omnibus the series matrix of samples analyzed using Agilent Human 1A Oligo UNC custom Microarrays (GPL1390; https://ftp.ncbi.nlm.nih.gov/geo/series/GSE26nnn/GSE26338/matrix/GSE26338-GPL1390_series_matrix.txt.gz) and used data as is. Differentially expressed genes were identified using the Significance Analysis of Microarray algorithm coded in the *samr* R package (Tusher et al. 2001). In SAM, we estimated the percentage of false positive predictions (i.e., False Discovery Rate, FDR) with 1,000 permutations and identified as differentially expressed those genes with FDR ≤ 5% and absolute fold change larger than a selected threshold (e.g. ≥ 2) in the comparison of primary tumors and metastases, with either paired and unpaired response types.

#### Survival analysis

Kaplain-Meier were generated with GOBO online tool (http://co.bmc.lu.se/gobo/gsa.pl) which calculates log-rank P value (Mantel–Cox method). EphB6 activity signature has been generated by selecting the most upregulated genes in coculture in cells with EphB6 knock-down (Supplementary Table S4), i.e. genes that are anti-correlated with EphB6.

### Immunofluorescence and imaging

#### Lysosomes visualisation

Cells were plated onto coverslips in MLNL medium and incubated with 50 nM LysoTracker Red DND-99 (ThermoFisher, L7528) for 30 minutes at 37°C prior to fixation (4% PFA for 12 minutes at room temperature and washed three times in PBS). Coverslips were mounted with ProLong Diamond Antifade Mountant with DAPI (Invitrogen, P36962). For quantification, at least 20 fields were acquired for each condition using the same acquisition settings. Images were analyzed with Fiji software. Percentage of Lysotracker^+^ cytoplasmic area was calculated according to the formula: Lysotracker^+^ area/(total cell area-nuclear area)* 100. Lysotracker^+^ area was determined with “Analyze particles” tool applying the same threshold for all the images.

#### Visualisation of mouse lungs with DDCCs

5 × 10^5^ D2.0R-EGFP expressing shCtrl, shEphB6#31 or shEphB6#35 cells were injected as indicated above (three mice/cell line). After 4 days, mice were culled and left ventricle perfused with 4% PFA to ensure optimal fixation of inner lung tissue. Lungs were then excised, fixed for 3 hours in 4% PFA and immersed in 30% sucrose for 72 hours. After incubation, lungs were embedded in O.C.T. compound (Histo-Line Laboratories, R0030) for rapid freezing with liquid nitrogen vapor. Frozen material was cut in 10 μm sections, fixed in 4% PFA for 10 minutes at room temperature and, after washes, permeabilized for 15 minutes in 0.2% Triton-X 100 in PBS. Blocking step was performed O/N at 4°C with 3% BSA, 0.02% Tween-20 in PBS. Primary and secondary antibodies were incubated in blocking buffer at room temperature for 4 and 1 hours, respectively, in a wet chamber; phalloidin was also added to secondary antibodies. The following antibodies and stain were used: Chicken anti-GFP (Abcam, ab13970, 1:200); Rabbit anti-Ki76 (Spring Bioscience Corp, M3062, 1:100); Alexa Fluor 488 Goat anti-chicken (Life technologies, A11039, 1:200); Alexa Fluor 568 goat anti-rabbit (Life Technologies, cat. A11036, 1:300); Alexa Fluor 647 phalloidin (Life Technologies, A22287, 1:100); Alexa Fluor 647 anti-LAMP2 (Santa Cruz Biotechnology, clone H4B4, sc-18822 AF647, 1:30); Hoechst (Sigma-Aldrich, B1155, 10 μg/ml). Slides were mounted with ProLong Diamond Antifade Mountant with DAPI (Invitrogen, P36962). Images of cells and lungs were acquired with Leica TCS SP8 MP confocal microscope employing the LasX software (40x objective). Ki67^+^ cells in contact with GFP^+^ cells were manually counted.

### Reporter assay

D2.0R (parental, EGFP-shCtrl and EGFP-shEphB6 #31 and #35) cells were transfected with Lipofectamine 3000 Transfection Reagent (Invitrogen, L3000001) following the manufacturer’s instructions. TFEB transcriptional reporter plasmid (RAGD promoter cloned upstream of luciferase gene, gift from Prof. Graziano Martello, University of Padua)(Malta et al. 2017) were transfected together with a plasmid with constitutive expression of Renilla luciferase to normalize for transfection efficiency (Montagner et al. 2016). After 6h, 1.8×10^4^ transfected cells were plated both on TT1 layer (coculture) and on plastic (monoculture) in 24-well format. 48h after replating, cells were harvested in Luc lysis buffer (25 mM Tris pH 7.8, 2.5 mM EDTA, 10% glycerol, 1% NP-40) and samples on plastic were diluted 1:5 in Luc lysis buffer to balance the Luciferase/Renilla content compared to coculture. Luciferase and Renilla activity were determined in a Tecan plate luminometer with freshly reconstituted assay reagents (0.5 mM D-Luciferin (Sigma-Aldrich, L9504), 20 mM tricine, 1 mM (MgCO_3_)_4_Mg(OH)2, 2.7 mM MgSO_4_, 0.1 mM EDTA, 33 mM DTT, 0.27 mM CoA, 0.53 mM ATP for Luciferase reaction, and 4 μg/ml coelenterazine (Invitrogen, C2944) in TBS 1X for Renilla reaction). Each sample was transfected at least in three biological duplicates in each experiment.

### Statistical methodology

Normal distribution of data was tested with Shapiro-Wilk test for experiments with sample size greater than ten. For sample sizes lower than ten, it is not easy to assess the underlying data distribution and non-parametric tests were preferred. For samples sizes lower than five, we preferred parametric tests owing to the minimum possible P value becoming large in the non-parametric case. For normally distributed samples, we performed Student’s two-tailed t-test for single comparisons (paired or unpaired) and ANOVA test (one-way or two-ways) for multiple comparisons. For non-normal data, we performed two-tailed Mann-Whitney test for single comparisons and Dunn’s test for multiple comparisons. Statistical analyses were performed with GraphPad Prism Software. Gene expression derived from microarray data of clinical samples was analysed with Significant Analysis of Microarray method (SAM, see Bioinformatics section). For survival plots (Kaplan–Meier analysis), data were analysed with GOBO (http://co.bmc.lu.se/gobo/gsa.pl) online tool which calculates log-rank P value (Mantel–Cox method). GSEA is generated from the GSEA online tool (http://software.broadinstitute.org/gsea/index.jsp), which also calculates the two primary statistics of the analysis: NES and false discovery rate (FDR). NES is calculated by normalizing enrichment score to gene-set size; FDR represents an estimated likelihood that a gene set with a given NES represents a false positive.

## Data availability

RNAseq data have been deposited at GEO Database and will be available concomitant with publication. Read-only access token is available to reviewers upon request. Other data that support the findings are available upon reasonable request from the corresponding authors.

## Acknowledgment

We are grateful to Julian Downward (Crick Institute), Dalit Barkan (University of Haifa) and Prof. R. Gomis (IRB, Barcelona) for gifts of cell lines. We are indebted to Sirio Dupont, and Graziano Martello (University of Padua) for thoughtful discussion and reagents. We are indebted to Flow cytometry, Experimental Histopathology, Bioinformatics and Biostatistics, Biological research, Cell services and Advanced sequencing facilities at the Crick Institute for exceptional scientific and technical support throughout the project. We thank Dr. Mattia Arboit for support with data visualization in R. C.R., S.H., & E.S. are supported by the Francis Crick Institute, which receives its core funding from Cancer Research UK (FC010144), the UK Medical Research Council (FC0010144) and the Wellcome Trust (FC010144). M.M. received funding from Marie Curie Actions—Intra-European Fellowships #625496 and BIRD Seed grant from Department of Molecular Medicine (University of Padua). C.D.H.R. was supported by a postdoctoral training award from Fonds de Recherche du Québec – Santé.

## Author contributions

M.Z.: conceptualisation, methodology, data curation, writing - review & editing. P. R: methodology. M.F., M.D., S.B.: formal analysis and visualisation of microarray data, writing - review & editing. P.C.: formal analysis and visualization of RNA sequencing data. S.D.: conceptualisation, funding acquisition, resources, writing - review & editing. C. D. H. R, S. H.: investigation, writing - review & editing. E.S, M.M.: conceptualization, formal analysis, funding acquisition, investigation, methodology, project administration, resources, supervision, visualization, writing - original draft.

---

**Supplementary Figure S1.**
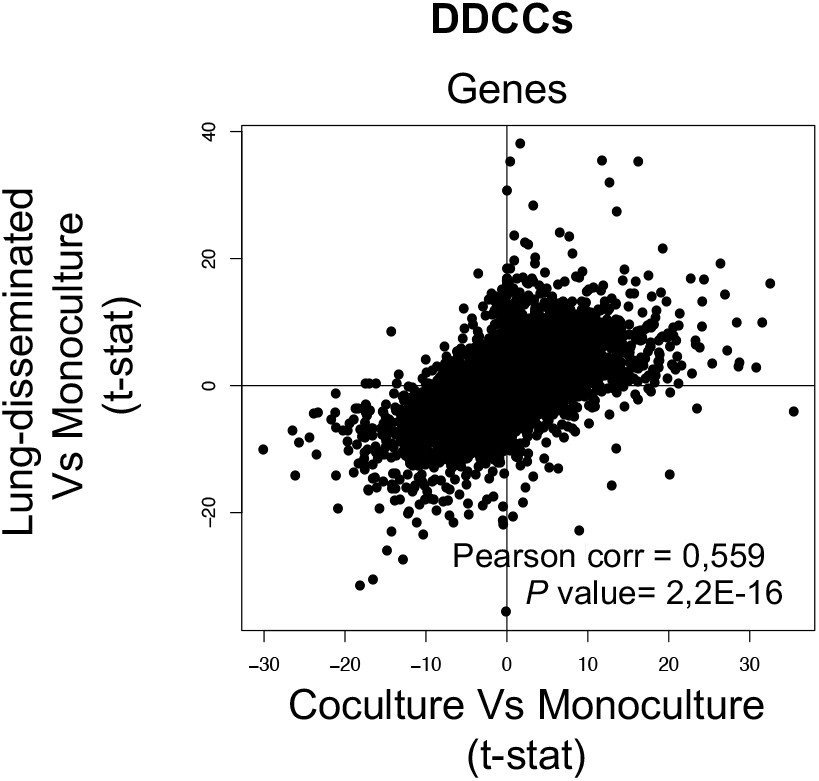
Scatterplot shows the correlation between the Wald’s statistic (gene level differences from DESEQ2) of D2.0R-EGFP cells disseminated in vivo (lungs), in coculture or monoculture as indicated.

**Supplementary Figure S2.**
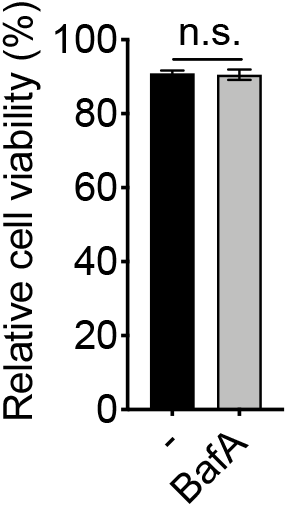
Cell viability, assayed by resazurin staining, of AT1-like cells treated or not with 4 μM Bafilomycin A for 4 days.

